# Sustained replication of synthetic canine distemper virus defective genomes *in vitro* and *in vivo*

**DOI:** 10.1101/2021.06.11.448162

**Authors:** Natasha L. Tilston-Lunel, Stephen R. Welch, Sham Nambulli, Rory D. de Vries, Gregory W Ho, David Wentworth, Reed Shabman, Stuart T. Nichol, Christina F. Spiropoulou, Rik L. de Swart, Linda J. Rennick, W. Paul Duprex

## Abstract

Defective interfering (DI) genomes restrict viral replication and induce type-I interferon. Since DI genomes have been proposed as vaccine adjuvants or therapeutic antiviral agents, it is important to understand their generation, delineate their mechanism of action, develop robust production capacities, assess their safety and *in vivo* longevity and determine their long-term effects. To address this, we generated a recombinant (r) canine distemper virus (CDV) from an entirely synthetic molecular clone designed using the genomic sequence from a clinical isolate obtained from a free-ranging raccoon with distemper. rCDV was serially passaged *in vitro* to identify DI genomes that naturally arise during rCDV replication. Defective genomes were identified by Sanger and next-generation sequencing techniques and predominant genomes were synthetically generated and cloned into T7-driven plasmids. Fully encapsidated DI particles (DIPs) were then generated using a rationally attenuated rCDV as a producer virus to drive DI genome replication. We demonstrate these DIPs interfere with rCDV replication in a dose-dependent manner *in vitro*. Finally, we show sustained replication of a fluorescent DIP in experimentally infected ferrets over a period of 14 days. Most importantly, DIPs were isolated from the lymphoid tissues which are a major site of CDV replication. Our established pipeline for detection, generation and assaying DIPs is transferable to highly pathogenic paramyxoviruses and will allow qualitative and quantitative assessment of the therapeutic effects of DIP administration on disease outcome.

**Importance:** Defective interfering (DI) genomes have long been considered inconvenient artifacts that suppressed viral replication *in vitro*. However, advances in sequencing technologies have led to DI genomes being identified in clinical samples, implicating them in disease progression and outcome. It has been suggested that DI genomes could be harnessed therapeutically. Negative strand RNA virus research has provided a rich pool of natural DI genomes over many years and they are probably the best understood *in vitro*. Here, we demonstrate identification, synthesis, production and experimental inoculation of novel CDV DI genomes in highly susceptible ferrets. These results provide important evidence that rationally designed and packaged DI genomes can survive the course of a wild-type virus infection.

## Introduction

Negative-sense (-) RNA viruses are prone to replication errors due to their low fidelity RNA-dependent RNA polymerase (RdRp). This results in a rich population of genetic variants, which may include subpopulations of defective interfering (DI) genomes. DI genomes are defined by their ability to disrupt standard genome replication, either by directly competing for resources or indirectly by triggering the interferon (IFN) pathway^1–5^. The most common and well-defined DI genomes include the deletion and copyback types^2^. During standard genome replication, the incoming (-) RNA is encapsidated by the viral nucleocapsid protein and contains a single genomic promoter (GP). RdRp binds to this GP and synthesizes complementary (+) RNA intermediates or antigenomes. The antigenomic promoter (AGP) present on the antigenomic RNA allows RdRp to bind and synthesize novel (-) RNA genomes^6^. However, when the RdRp prematurely dissociates from either its (-) or (+) RNA template it can reinitiate replication by binding (a) upstream of the same template and generate deletion genomes, or (b) onto the nascent complementary strand and generate ambisense genomes known as copybacks. Deletion and copyback genomes that retain functional replication and packaging signals can be maintained alongside the standard genome by utilizing it as a source for missing proteins. This association results in a predator-prey type scenario which can be visualized as cyclic titer patterns during *in vitro* passage experiments^2,7^.

The ability of DI genomes to disrupt virus replication has led to propositions for their use as vaccine adjuvants or antivirals^8–10^. Various studies using vesicular stomatitis virus (VSV)^11,12^, Sendai virus (SeV)^13^, human respiratory syncytial virus (HRSV)^14,15^ and influenza virus^16–18^ have demonstrated reduced standard viral yields both *in vitro* and *in vivo* when DI genomes are present. DI genomes are highly immunostimulatory in nature and demonstrate preferential interaction with RIG-I over the standard genome^19^. Copyback DI genomes specifically have been shown to stimulate production of several proinflammatory cytokines and chemokines, and enhance dendritic cell maturation^14,20–22^. In humans, DI genomes detected in HRSV positive samples correlate with increased expression levels of antiviral genes^23^, while their absence in influenza A virus (IAV) infected patients correlate with disease severity^24^. The antiviral activity of DI genomes has been assessed using both unencapsidated “naked” DI RNA, as well as DI-RNA packaged in a ribonucleoprotein complex, defined as a DI particle (DIP). In immunization studies, inactivated viruses adjuvanted with DI RNAs score better than controls with poly-IC or alum, by inducing type-I humoral and cellular immune responses and enhancing antibody levels^22,25^. This type-I IFN inducing ability allows DIPs to also protect against heterologous virus infection, as seen with influenza A virus (IAV) DIP 244/PR8 which protects mice from unrelated pneumonia virus^26^. However, for a DIP-based therapeutic to work dosage is vital. For instance, mice treated with 400 hemagglutinating units (HAU; 1.2 µg) of IAV DIP 244/PR8 one week prior to a second IAV challenge were completely protected, whereas mice that were treated with a ten-fold higher dose did not have the same outcome^16^. These results illustrate how achieving a potent dosage optimal to both outcompete standard virus replication and to induce an immune response is complex.

Another challenge when considering DIPs is their production. DIPs require appropriate packaging to allow successful delivery of the DI genome of sufficient potency to the right target cells. DI genomes by their very nature are replication-deficient and thus need a replication-competent helper virus to drive their production. The inherent enigma here is that the DI genome will also interfere with replication of the helper virus, thereby impeding a straightforward production process. The second challenge is that the DIPs need to then be purified from the helper virus. Certain viruses like SeV^13^, VSV^11,12^ and HRSV^27^ exhibit a virion size difference depending on whether the full-length or DI genome is packaged. This allows DIPs to be purified via density ultracentrifugation. This size difference is not seen in all viruses, nor would separation on basis of size be advisable for highly pathogenic viruses. Another option is ultraviolet (UV) irradiation, which is used to exploit the difference in genome length between the helper virus and DIP. At a precise dosage the much larger full-length genome can be selectively inactivated whilst leaving the smaller DI genome intact^28^. However, practically this is not a viable option when dealing with highly pathogenic BSL-4 pathogens, for example Nipah virus. Recently, a packaging cell-line expressing the missing IAV protein PB1 demonstrated successful production of pure IAV DIP 244/PR8 without the need for a helper virus^29^. This is an ideal scenario, as establishing such packaging cell-lines expressing the repertoire of proteins required for many of the other viral DIPs may not be straightforward.

In this study, we investigated a novel, rationally attenuated, recombinant (r) canine distemper virus (CDV) as a “producer virus” (i.e., helper virus), in order to generate rCDV DIPs. CDV is a morbillivirus in the paramyxovirus family, and similar to many other members it requires a polyhexameric genome length for replication^30^. Therefore, we focused on rule-of-six compliant rCDV DI genomes that were generated naturally (n) during *in vitro* passages. We developed assays to generate these rCDV DIPs using our producer virus system and assessed the DIPs for interference activity. Most importantly, we demonstrate that a synthetically (s) engineered fluorescent defective genome can successfully replicate and be maintained in ferrets during the course of a natural rCDV infection. CDV is a tractable biosafety level (BSL)-2 pathogen and ferrets are a naturally susceptible animal model, making our system a good resource for developing highly complex assay systems such as the ones required for DIPs.

## Results

### A single dominant defective viral genome generated early during rCDV^RI^ *in vitro* infection is consistently maintained across subsequent passages

To identify predominant DI genomes arising naturally during a CDV infection, we serially passaged four different versions of recombinant (r) CDV strain Rhode Island (rCDV^RI^) 10 times in Vero cells modified to express canine (c) CD150 (Vero-cCD150). First, a plasmid encoding the full-length rCDV^RI^ antigenome was modified to encode rCDVs expressing reporter proteins Venus, monomeric blue fluorescent protein (TagBFP), dTomato fluorescent protein (dTom) or *Gaussia luciferase* (Gluc), from an additional transcription unit (ATU) at position six in the genome. Viruses rCDV^RI^Venus(6), rCDV^RI^TagBFP(6), rCDV^RI^dTom(6) and rCDV^RI^Gluc(6) were generated in Vero-cCD150. Here, cells were first infected with a recombinant vaccinia virus expressing T7 polymerase (MVA-T7) and then transfected with expression plasmids expressing the nucleocapsid (N), phospho-(P) and large (L) proteins along with the full-length plasmids pCDV^RI^Venus(6), pCDV^RI^TagBFP(6), pCDV^RI^dTom(6) or pCDV^RI^Gluc(6). Virus was rescued 5 – 7 days post-transfection (d.p.t.), and a clonal population was generated by plaque picking syncytia.

In the first experiment, rCDV^RI^Venus(6) was passaged once (P1) at MOI 0.05 in Vero-cCD150 cells. This stock was then serially passaged nine times, in triplicate (A, B and C) (Figure 1A). We found that passages A, B and C followed a highly similar viral titer pattern over the 10 passages (Figure 1B), which we expect is due to identical DI genomes in all three experiments. Using a copyback genome-specific RT/PCR assay ^31^ (Table 1, priCDV^RI^-A1R and priCDV^RI^-A2R) we amplified a 720 nt copyback genome in passage B at P10 (Figure 1C). To determine if a similar outcome would arise when repeated, rCDV^RI^TagBFP(6), rCDV^RI^dTom(6) and rCDV^RI^Gluc(6) were serially passaged under similar conditions to rCDV^RI^Venus(6) (Figure 1D). This time, we observed a different titer pattern for each virus (Figure 1E). rCDV^RI^TagBFP(6) and rCDV^RI^Gluc(6) titers crashed at passage 3, with rCDV^RI^Gluc(6) crashing also at P6 and P10. Viral titers for rCDV^RI^dTom(6) on the other hand appeared to remain relatively constant throughout the 10 passages. Here, RT/PCR results revealed a unique copyback genome for each virus, 630 nt in rCDV^RI^TagBFP(6), 1092 nt in rCDV^RI^dTom(6) and 690 nt in rCDV^RI^Gluc(6) (Figure 1F). Genome lengths of the copyback sequences identified by RT/PCR from these passage experiments are compliant with the rule-of-six requirement for morbilliviruses.

**Figure 1.**
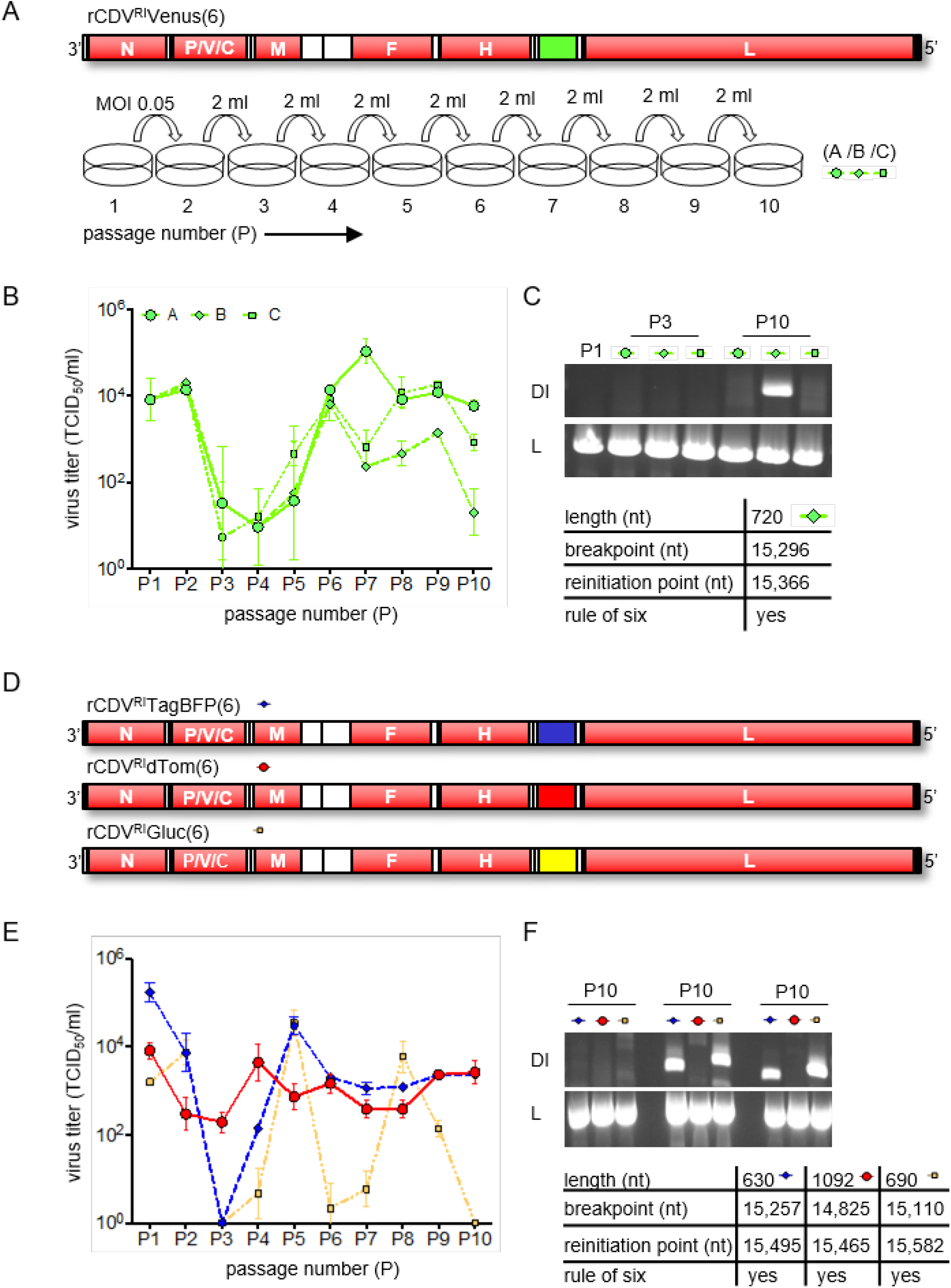
Serial in vitro passage of rCDVRI. (A) rCDVRIVenus(6) was rescued, plaque picked and passaged once (P1) in Vero-cCD150 cells. Nine serial passages were carried out in a fixed volume (2 ml) in triplicate (A, B and C). (B) samples from each passage were titrated in Vero-cCD150 cells and represented as TCID50/ml. Error bars represent standard deviation (n=3). (C) RT/PCR was carried out on passages A, B and C. cbgenome-specific primers detected nDIRI07cb (length 720 nt) in B at P10. Part of L was amplified as a positive control for viral RNA. (D) rCDVRITagBFP(6), rCDVRIdTom(6) and rCDVRIGluc(6) were rescued, plaque picked and serially passaged 10 times in a fixed volume (2 ml) in Vero-cCD150 cells. (E) viral titers were determined as above. (F) RT/PCR using cbgenome-specific primers amplified different DI genomes for rCDVRITagBFP(6), rCDVRIdTom(6) and rCDVRIGluc(6) corresponding to lengths 630 (nDIRI04cb), 690 (nDIRI11cb) and 1092 (nDIRI12cb) nts respectively. Viral RNA was confirmed in all samples by amplifying L.

Next, all passaged samples were sequenced using the Illumina MiSeq platform. Datasets were first quality filtered and aligned to their respective reference genome using CLC Genomics Workbench. Datasets were then examined to identify chimeric reads, which either consist of deletion junctions or head-to-tail rearrangements of the genome. These chimeric points were subsequently mapped onto full-length reference genomes, and break and reinitiation points (i.e., the sequence where the RdRp dissociates from the RNA template and the sequence where the RdRp reinitiates replication) were identified. In the final dataset, to help eliminate false positives that may have passed through the initial screening, we applied a read cutoff value of 1. Using this dataset, the frequencies of every defective genome identified across the passages for each virus were plotted. This revealed that each passage contained one predominant defective genome (Figure 2A and B). Mapping information identified these copyback genomes identical to those amplified by RT/PCR, hence providing confidence in our chimeric Illumina dataset.

**Figure 2.**
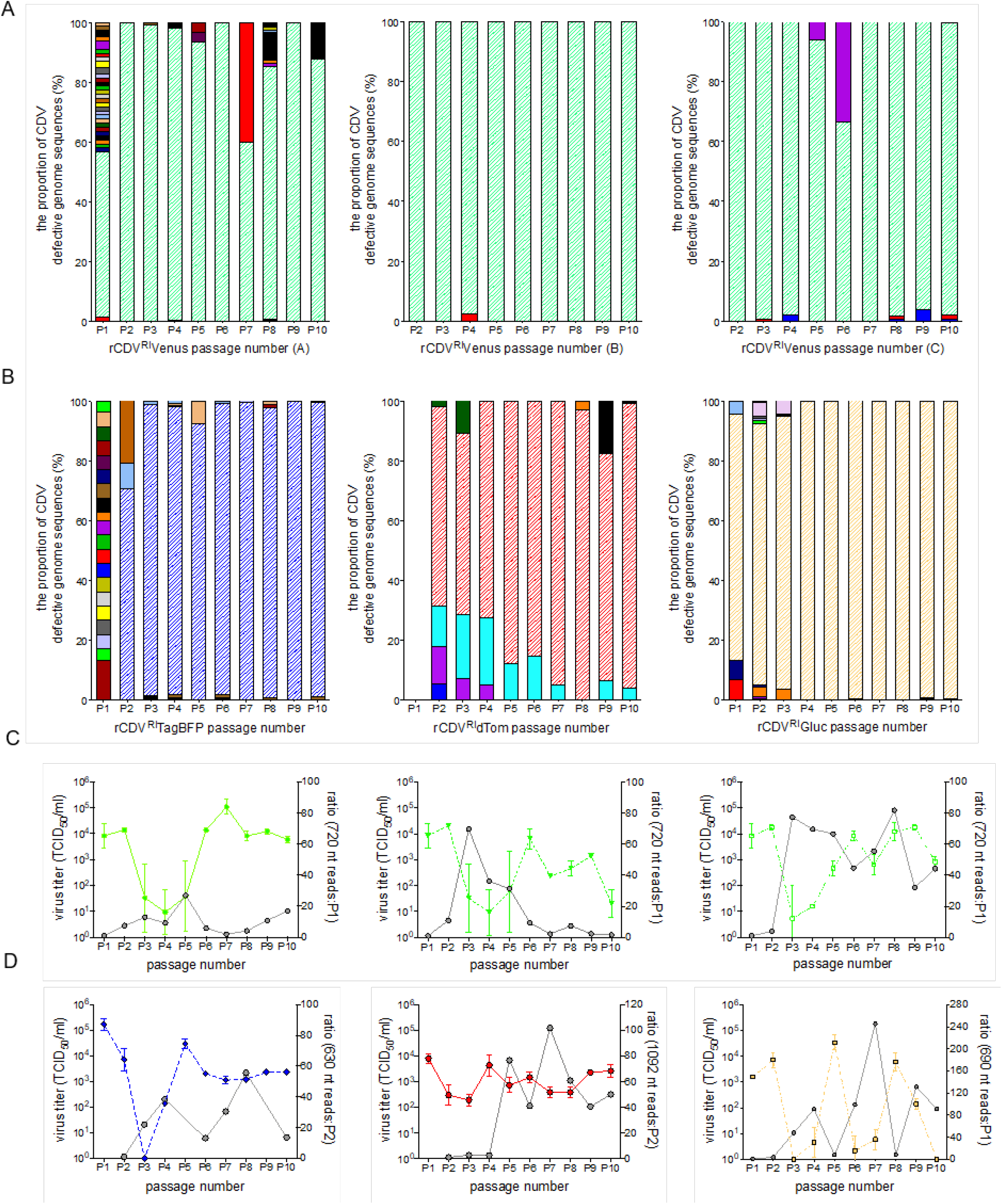
rCDV^RI^ defective genomes at each passage. (A) and (B) the relative frequency of every CDV defective genome at the 10 passages. Each genome is represented by a different color. A read cut-off value of 1 was applied to eliminate sequence noise. (C) and (D) sequence reads for the predominant DI genome from (A) and (B) overlaid onto the viral titers from their corresponding passage experiment. Ratio of reads were calculated from when the predominant DI genome first appeared in passage. i.e., the diagonal striped areas in (A) and (B) represent the predominant DI genome.

In the first experiment using rCDV^RI^Venus(6), a 720 nt genome was identified at P1, and even though passage A, B and C were independent from P2 onwards, the 720 nt genome remained predominant throughout the nine passages (Figure 2A). In the second experiment, a 630 nt genome was identified at P2 in rCDV^RI^TagBFP(6) and remained predominant till P10 (Figure 2B). Whether this genome formed at P1, but was just undetected is unclear. In rCDV^RI^dTom(6), no defective genomes were detected at P1, from P2 onwards a 1092 nt copyback genome became predominant. With rCDV^RI^Gluc(6) a 690 nt copyback genome prevailed from P1 up until P10 (Figure 2B). A ratio of the 720 nt, 630 nt, 690 nt and 1092 nt copyback genome sequencing reads was calculated for each passage based on when they were first detected. Overlaying these numbers onto their respective parental virus titers demonstrated an inverse correlation in cyclic patterns of DI genome reads and virus titer (Figure 2C and D). This was especially convincing for rCDV^RI^Gluc(6) (Figure 2D).

In brief, we identified 15 rule-of-six compliant rCDV copyback genomes from the serial passages and an additional six during the course of this study (Table 2). We found only four copyback genomes that would have formed towards the GP end of the genome (i.e., GP copybacks). Breakpoints for these GP copyback genomes fell in the N and F gene, and were not rule-of-six compliant. Plotting all AG copyback break and reinitiation points, whether or not these genomes were rule-of-six compliant, we found that breakpoints in our dataset typically fell between nts 13,371 and 15,624 of the rCDV^RI^ genome, and clustered between nt 15,250 and 15,450 (L gene). Reinitiation points for these genomes as expected occurred much closer to the genome end, between nt 15,300 and 15,683 (Figure 3A). We found two snap-back genomes (with identical breakpoint and reinitiation sites), one of which was rule-of-six compliant (2124 nt, Table 2). In terms of defective genome uniqueness, we found more rule-of-six deletion genomes than copyback genomes (Figure 3B). However, a majority of these fell below our read cut-off threshold of 1. It is unclear at this time whether these genomes are sequencing artifacts or true deletion sequences that were simply unable to compete with the predominant copyback genome.

**Figure 3.**
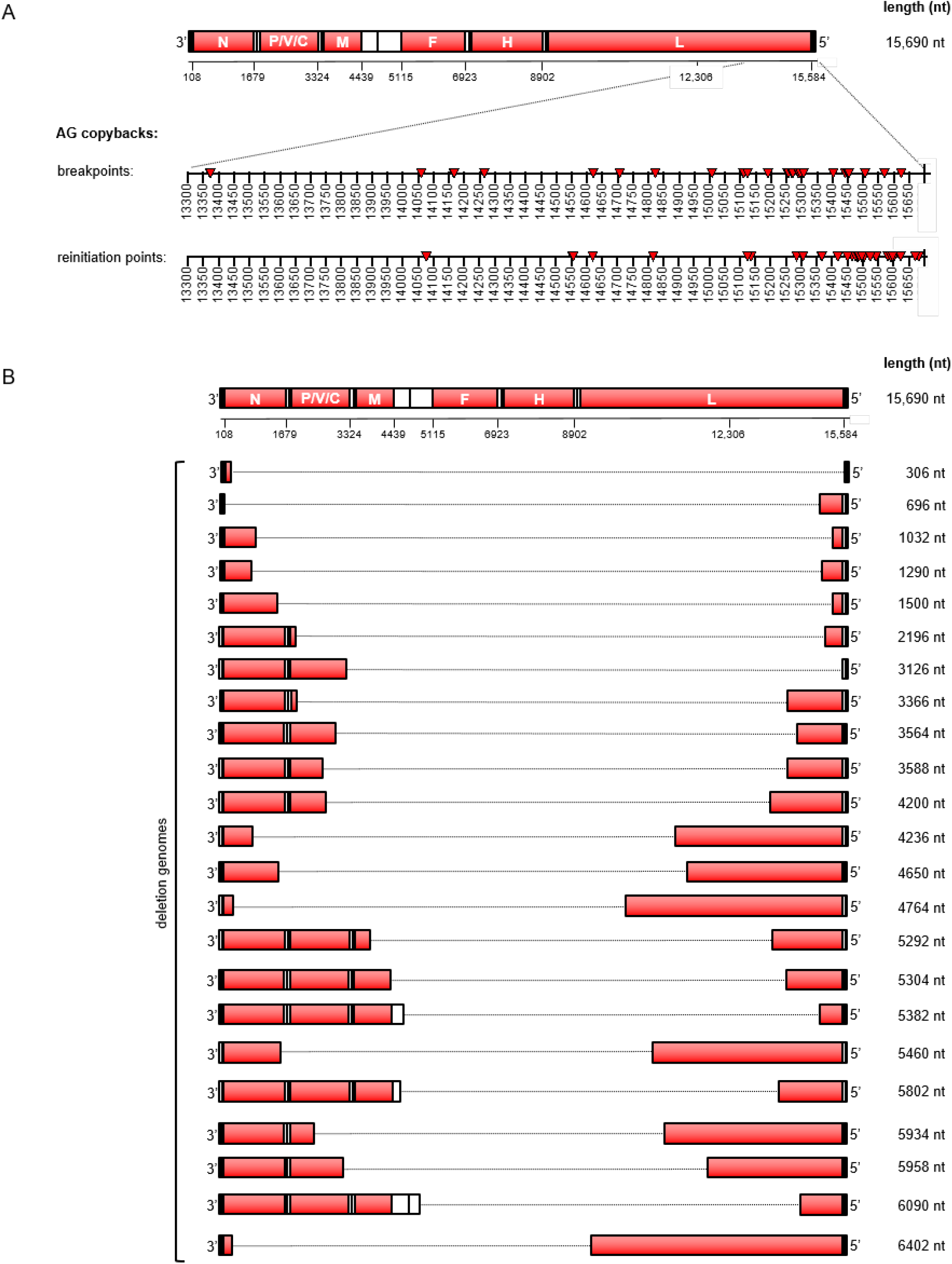
rCDV^RI^ defective genome break and reinitiation points. (A) breakpoints and reinitiation points for AG copyback genomes identified in experiments (B) and (E) are plotted onto a scale illustrating their location on rCDV^RI^. (B). All rule-of-six compliant deletion genomes identified in experiments (B) and (E).

### Generation of DIPs using a rationally attenuated rCDV^RI^ as a producer virus

To produce DIPs in a non-pathogenic background, we generated an attenuated rCDV^RI^Venus(6), using an approach as described in our previous work^32,33^. We predicted the second variable hinge in CDV^RI^ L gene to be between nucleotides 14,111 and 14,180. We mutated the genome positions 14,826-27 and 14,835-36 in plasmid pCDV^RI^Venus(6) from ‘CT’ to GG and AA, respectively. This created restriction sites *Msc* I and *Hpa* I, which were used to clone the open reading frame of enhanced green fluorescent protein (EGFP) into the L gene (Figure 4A). rCDV^RI^Venus(6)-LEGFP was rescued and plaque picked (P0) in Vero-cCD150 cells, with titers similar to rCDV^RI^Venus(6). This virus thus encoded both Venus (from an ATU) and EGFP (fused to the viral polymerase). In previous studies we have demonstrated that this fusion protein remains functional as a polymerase, but with reduced efficacy thus explaining the viral attenuation phenotype.

**Figure 4.**
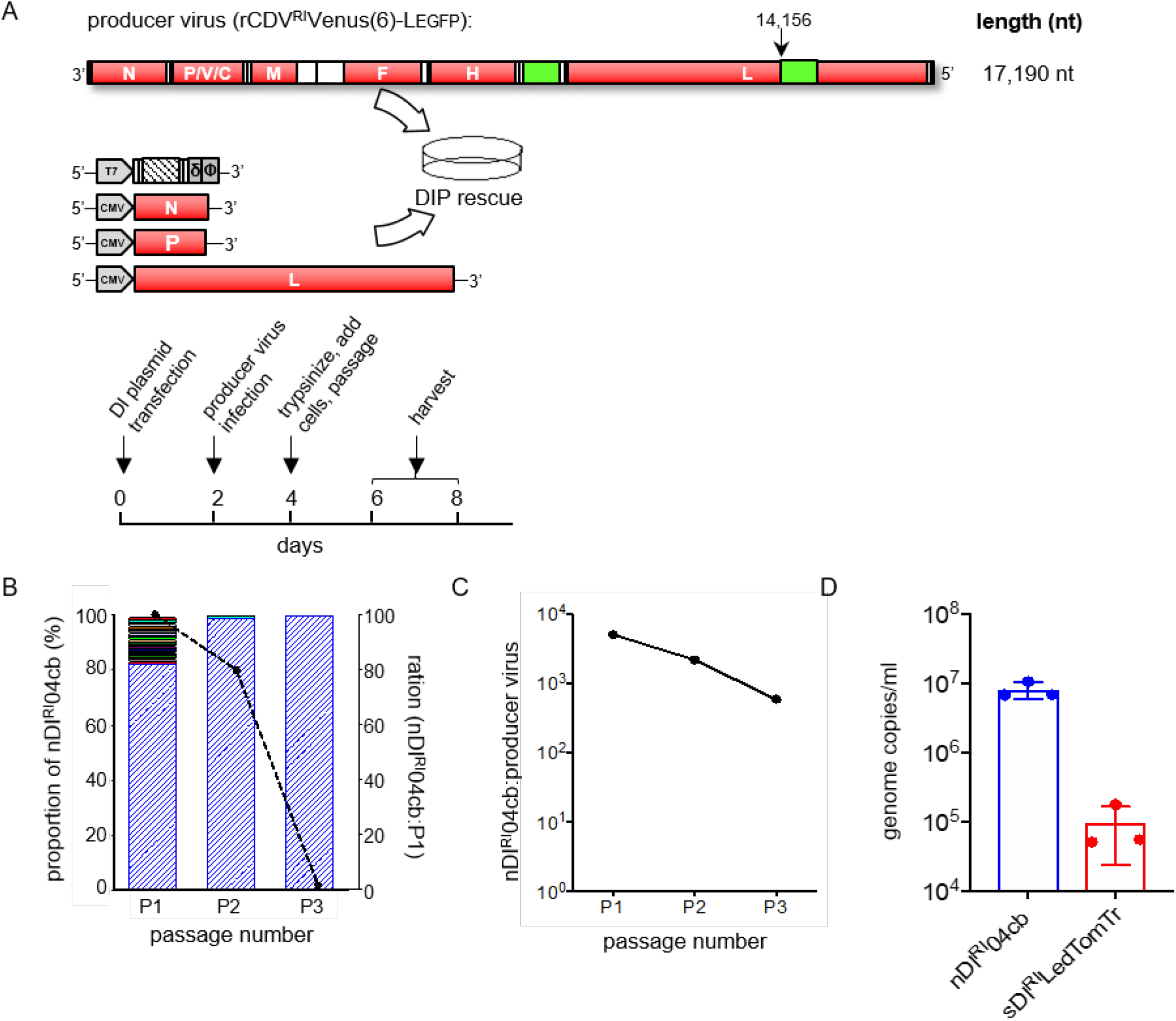
Production of rCDV^RI^ DIPs using a recombinant producer virus. (A) experimental set-up of an rCDV^RI^ DIP rescue. (B). frequency of rCDV^RI^ defective genomes generated as shown in (A) using nDI^RI^04cb DIP. Rescued DIP stock still including the producer virus was passaged three times in Vero-cCD150 cells. Blue bars represent the predominant nDI^RI^04cb. The overlaid line graph represents sequence reads for the three passages. (C). qRT-PCR for (B) shows the ratio of DI/full-length genome at three passages. (D). controlling DIP production. qRT-PCR for a passage 2 experiment where a determined ratio of DI/full-length was supplied from passage one. The experiment was performed in triplicate. Error bars represent standard deviation of titers (n=3).

Next, we investigated whether a clonal population of DIPs can be produced using rCDV^RI^Venus(6)-LEGFP. We first constructed DI genome-expressing plasmids by inserting synthetic copyback (cb) DI genome sequences for nDI^RI^04cb, nDI^RI^07cb, nDI^RI^10cb and nDI^RI^11cb (Table 2) into a T7-driven plasmid backbone. We also constructed a deletion (del) genome (sDI^RI^LedTomTr) by placing the dTom sequence between the leader and trailer sequences of rCDV^RI^ genome. To then generate DIPs, we transfected Vero-cCD150 cells (supplemented with T7 polymerase) with a DI-plasmid and CMV-driven helper plasmids expressing rCDV^RI^ N, P and L proteins. Two d.p.t. these cells were superinfected with rCDV^RI^Venus(6)-LEGFP at an MOI 0.001, and following an experimental setup described in Figure 4A stocks containing DIPs were produced.

To determine the defective genome population in rescued (P0) stocks, we passaged DIP nDI^RI^04cb an additional three times in Vero-cCD150 cells and sequenced them using our Illumina NGS pipeline. At P1 about 20% of the defective genome reads were non-rule-of-six copyback and deletion sequences, with only 1 sequencing read each (Figure 4B, left axis). These numbers decreased drastically at P2 and P3 leaving nDI^RI^04cb as the predominant sequence, although the overall number of reads in the samples also dropped (Figure 4B, right axis). This highlighted a production issue when using a producer virus, due to the competition between DI and full-length genome (Figure 4C). To solve this issue, we first quantified the genome copies of nDI^RI^04cb or sDI^RI^LedTomTr present in P1, and the number of full-length genome copies in our producer stock using a qRT/PCR assay (Table 1). We then empirically determined that a low DI:virus ratio (0.05:1) repeatedly generated between 10^5^ - 10^7^ copies/μl of DI genome (Figure 4D), providing a way to control DIP output.

### UV irradiation for targeted inactivation of producer virus

To assess defective genome-specific effects without the confounding effects of the producer virus, we first tested various UV dosages that would sufficiently inactivate only the full-length genome. Briefly, producer virus samples were irradiated using a UV crosslinker at UV dosages between 0 to 60 mJ/cm^2^, at 2 ml in a 6-well tray. By 40 mJ/cm^2^ no infectious virus could be detected via TCID_50_ (Figure 5A).

**Figure 5.**
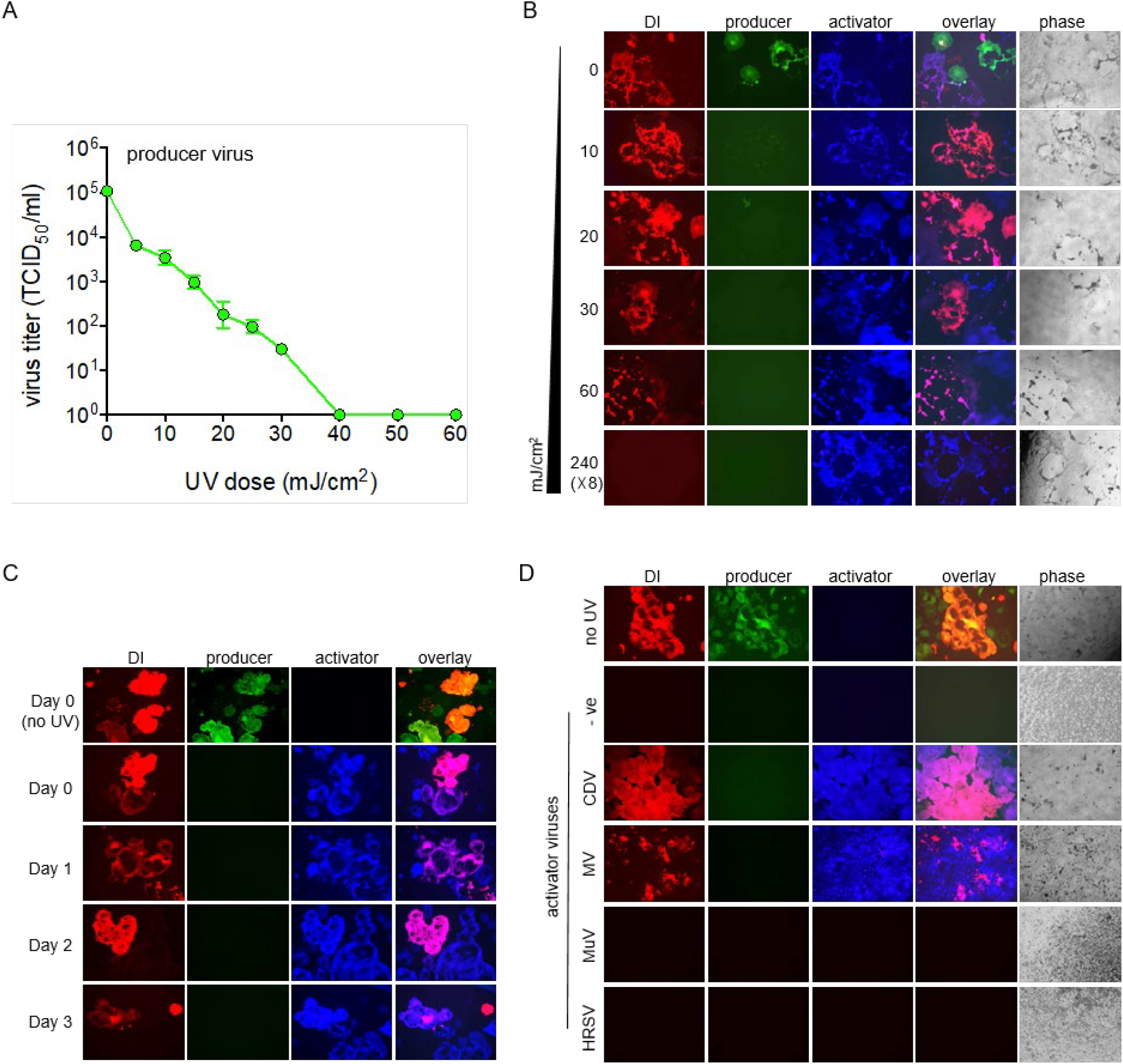
UV inactivation of “producer” virus and activation of DI with “activator” virus. (A) the effect of various UV dosages on rCDV^RI^ “producer” virus titers. Samples were irradiated using an UV crosslinker (CX-2000 Crosslinker, UVP) and TCID_50_/ml determined. (B) UV doses were tested in the presence of a DI genome (red) empirically to determine required amount to inactivate rCDV^RI^ producer virus and not the DI genome. Vero-cCD150 cells infected with UV-treated samples were superinfected with rCDV^RI^TagBFP “activator” virus, in order to drive the replication of the DI in the samples. (C) stability of the DI genome in infected cells were tested. DI genomes remained ‘dormant’ in cells for up to three days as shown by genome activation with rCDV^RI^. (D) activation of rCDV^RI^ DI genome using various paramyxoviruses as “activator” virus. Cross reactivity of rCDV^RI^ DIP was tested against other paramyxoviruses measles virus (MV), mumps virus (MuV) or human respiratory syncytial virus (HRSV). Activation of a rCDV^RI^ DI genome only occurred with CDV and MV, both morbilliviruses.

Next, we tested these UV dosages in our DIP stocks in order to determine if the DI genome remained active. Since an active full-length genome is required for defective genome replication, we superinfected Vero-cCD150 cells that were infected with a range of UV-treated DIP stocks. Here, we used rCDV^RI^TagBFP(6) as an ‘activator virus’ (blue syncytia, Figure 5B and C). We found that the defective genome (red syncytia, Figure 5B) remained active even at 60 mJ/cm^2^. In the subsequent experiments we therefore chose 60 mJ/cm^2^ for producer virus inactivation. For complete inactivation of DI-genome we used a high UV dose of 240(×8) mJ/cm^2^, as a single dose proved insufficient. Interestingly, we also found that defective genomes can remain inactive in cells for at least 3 days post infection until superinfected with an activator virus (Figure 5C).

Next, to confirm whether defective genome activation was specifically due to the activator virus and/or if other paramyxoviruses can replicate the defective genome, we infected Vero-cCD150 cells with DIPs and then attempted to activate expression using CDV, measles virus (MV), mumps virus (MuV) or HRSV. Activation of a CDV defective genome occurred with CDV and MV, but not with MuV or HRSV (Figure 5D).

### rCDV^RI^ DIP specific interference activity is dose dependent

We tested four cbDIPs: nDI^RI^04cb, nDI^RI^07cb, nDI^RI^10cb and nDI^RI^11cb (Table 2), and sDI^RI^LedTomTr for their ability to interfere with rCDV^RI^ replication *in vitro* using Vero-cCD150 cells which lack a fully functional IFN system. Vero-cCD150 cell monolayers infected with a 10-fold serial dilution of the DIP stocks demonstrated almost complete inhibition of rCDV^RI^ infection at the highest DIP concentration (Figure 6A). Using cb DIP nDI^RI^04cb, we determined if inhibition was DIP-specific or an effect due to cytokines and/or cell debris. Here, DIPs were UV-inactivated at 120 (×8) mJ/cm^2^ to ensure complete DI genome inactivation. Next, Vero-cCD150 cells were infected with various ratios of rCDV^RI^ and active DIPs or rCDV^RI^ and inactivated DIPs. Ratios were based on genome copies determined using a qRT/PCR assay. We found a dose-dependent DIP-specific interference affect, in which a genome ratio of 10:000 DIP:1 rCDV^RI^ was required to completely inhibit virus replication in Vero-cCD150 cells (Figure 6B). Importantly, we observed no reduction in rCDV^RI^ yield with the inactivated DIPs.

**Figure 6.**
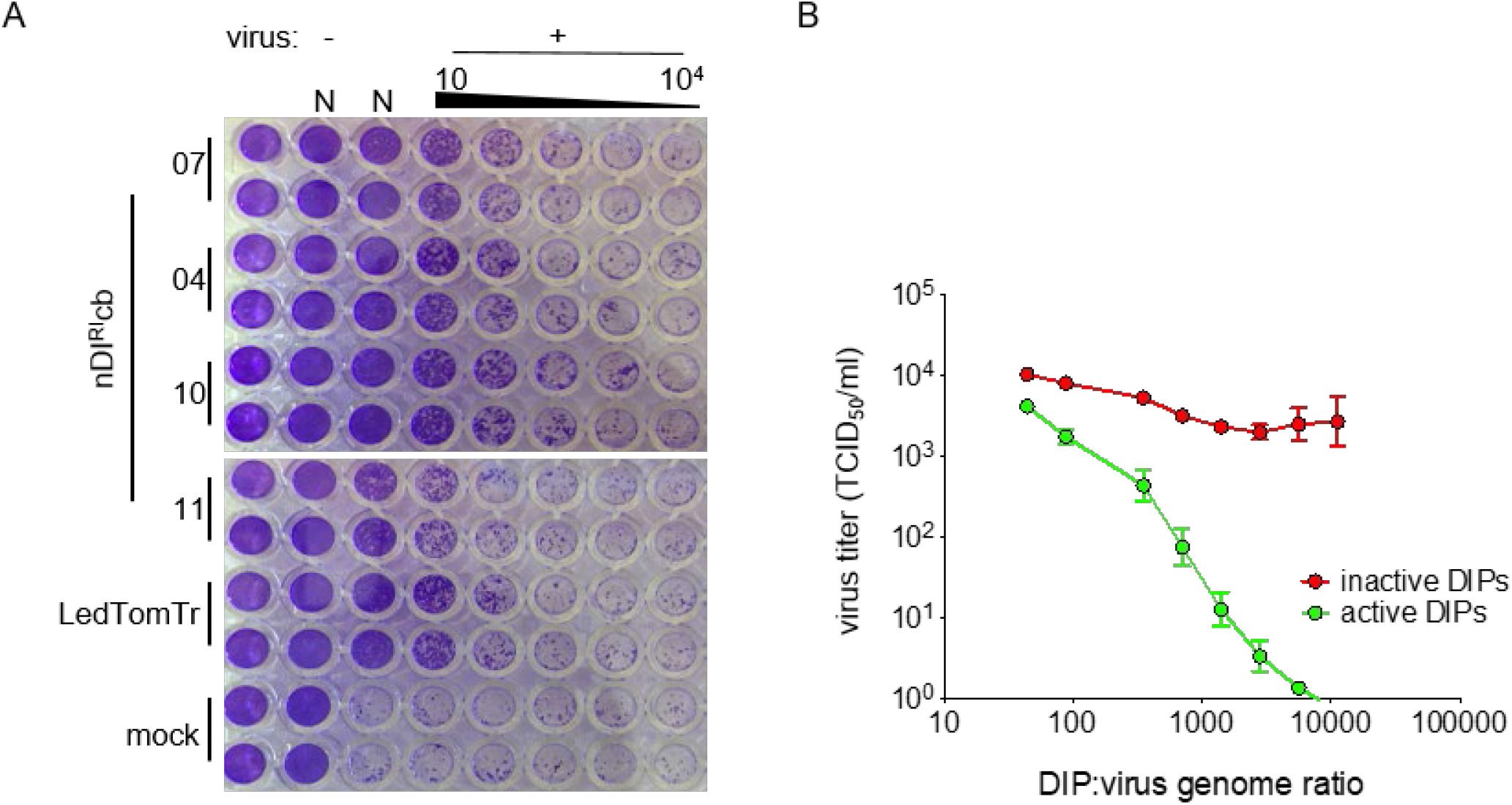
DIP interference on rCDV^RI^ infection *in vitro*. (A) copyback nDIs 07, 04, 10 and 11 (Table 2) and deletion DIP sDI^RI^LedTomTr were tested for their ability to interfere with rCDV^RI^ replication *in vitro*. The amount of inhibition to rCDV^RI^ infection in Vero-cCD150 cells is visualized by cytopathic effect (CPE) (cell monolayers stained by crystal violet. N=neat, undiluted DIP stock). (B) dose response curve for copyback nDI^RI^04cb. To obtain a dose at which a DIP completely eliminates CDV infection Vero-cCD150 cells were treated with specific ratios virus and DIPs (green line). Control infections were performed using UV-inactivated DIPs (red line) to demonstrate interfering effects were DI genome replication specific. rCDV^RI^ titers were measured by TCID_50_/ml at 72 h.p.i.

### rCDV^RI^ DIPs replicate in appropriate cells in a ferret model

To assess if DIP nDI^RI^04cb can replicate and be maintained during the course of infection in a natural host animal model. We first demonstrated that DIPs (delDIP expressing Venus) could successfully replicate in disease-relevant IFN-competent canine B-cell lymphoma cell line (CLBL-1) when co-infected with rCDV^RI^TagBFP(6) as an activator virus (Figure 7A). Next, four groups of three ferrets were infected with rCDV^RI^TagBFP(6) as an activator virus. Animals were either co-administered with nDI^RI^04cb or DMEM (control), or pre-administered with nDI^RI^04cb or sDI^RI^dTomdel04cb (nDI^RI^04cb variant expressing dTom). DIPs were pre-administered 6 hours prior to activator virus infection. All infections were carried out via the intra-tracheal (IT) route (Figure 7B). Animals were monitored over a 14-day period for clinical signs and symptoms, and blood samples were collected at 0 d.p.i. and then every 2 days. No weight loss was observed (Figure 7C) over the infection period. All animals had lymphopenia, (Figure 7D) which is typical in ferrets infected with CDV. White blood cells (WBC) were isolated from the blood samples and the percentage of TagBFP positive cells were determined by flow cytometry (Figure 7E), this was confirmed at the same time by virus isolation in Vero-cCD150 cells. rCDV^RI^TagBFP(6) was detected in WBC of all ferrets with a peak at 6 d.p.i. (Figure 7E). At necropsy, rCDV^RI^TagBFP(6) was also isolated from the lymph nodes of all animals. Importantly, we isolated sDI^RI^dTomdel04cb in two animals from WBC at 6 d.p.i. and lymph nodes at 14 d.p.i. (Figure 7F). Using a genome-specific and DI-specific qRT/PCR assay we detected low levels of DI genomes in the WBC and lymph nodes for both sDI^RI^dTomdel04cb (Figure 7G and H) and nDI^RI^04cb (Figure 7I and J).

**Figure 7.**
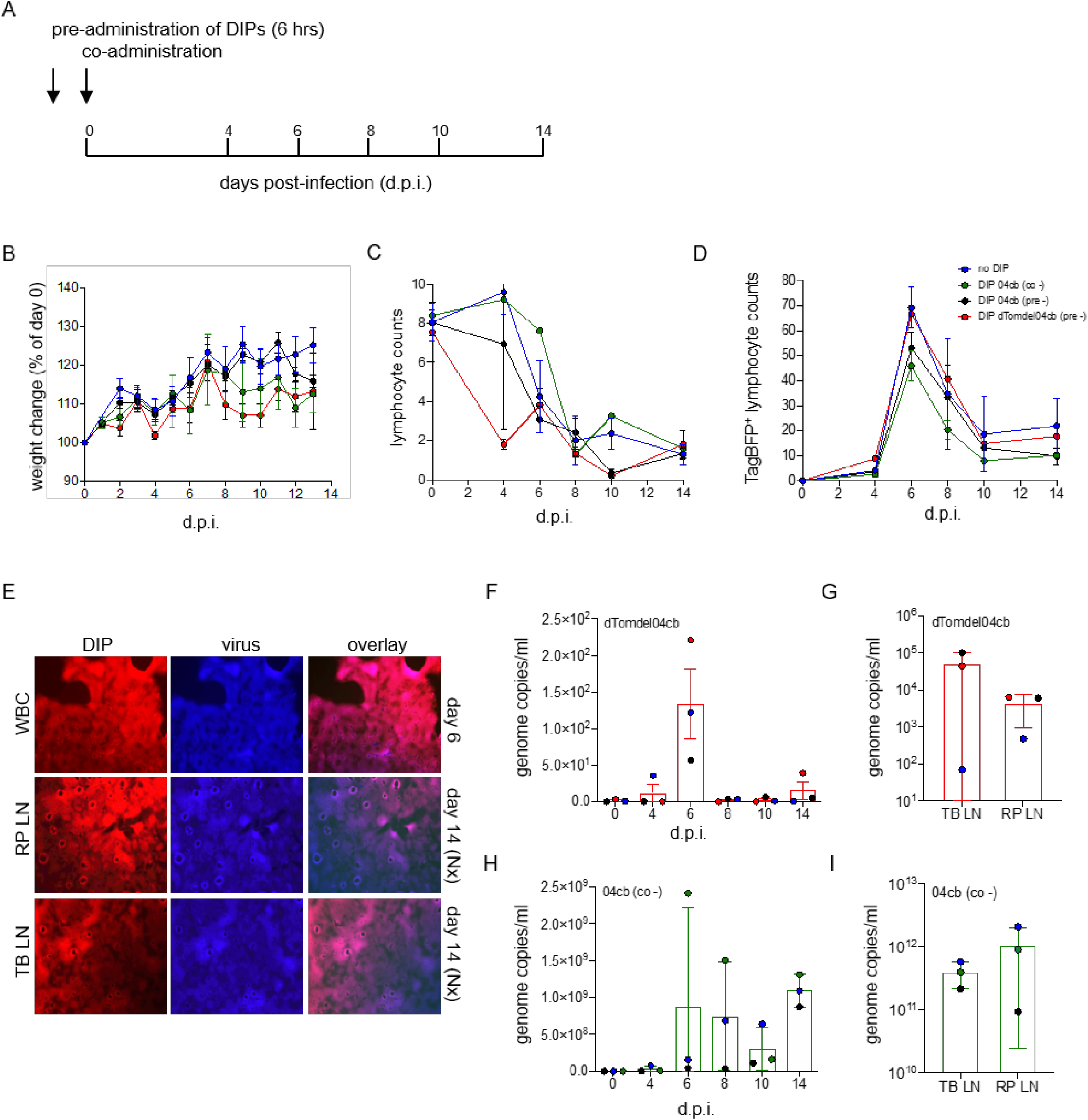
rCDV^RI^ infection in ferrets. (A) experimental outline of ferret infection. Ferrets were either pre- or co-infected with DIP nDI^RI^04cb, pre-inoculated with sDI^RI^dTomdel04cb, or culture medium prior to infection with the challenge virus rCDV^RI^TagBFP. Pre-administration was carried out 6 hours before infection.(B) weight change of ferrets during the course of infection. (C) lymphocyte counts in EDTA blood determined by flow cytometry shows lymphopenia. (D) virus infection in lymphocytes was determined for each time point by monitoring fluorescence using flow cytometry. Peak in infection is seen at day 6. (E) sDI^RI^dTom04cb and virus were isolated from WBC (6 d.p.i.) and from lymph nodes (14 d.p.i.) from ferrets. TB-LN, tracheol-bronchial lymph node; RP-LN, retropharyngeal lymph node. (F) and (G) sDI^RI^dTom04cb genome copies were detected in WBCs at each time point during the course of the infection and at necropsy in single cell suspensions of lymphoid tissues. (H) and (I) nDI^RI^04cb genome copies were detected in WBCs at each time point during the course of the infection and at necropsy in lymphoid tissues.

## Discussion

Viruses adapt quickly to their host environment and so it is crucial to minimize their passage *in vitro* if we are to obtain meaningful information. In the present study, we describe the establishment of a novel recombinant canine distemper virus strain Rhode Island (rCDV^RI^). rCDV^RI^ is the first non-laboratory adapted rCDV, based on a currently circulating wild-type strain isolated from a raccoon on Rhode Island, USA in 2012. We propose that rCDV^RI^ should replace all currently used laboratory-adapted rCDV strains such as Snyder-Hill and A75 in pathogenesis studies. The accessibility and affordability of sequencing and synthetic biology should now allow the use of field isolates of known provenance such as CDV^RI^ to replace laboratory-adapted strains.

The overarching goal of this study was to assess the ability of a non-laboratory adapted CDV to generate DI genomes *in vitro* and whether these DI genomes could be maintained *in vivo* in a natural animal model. We used a combination of RT/PCR/Sanger sequencing and short-read Illumina sequencing to identify defective genomes in rCDV^RI^. We used a bioinformatics pipeline^34^ to identify DI-junctions in the Illumina datasets, and found that all high frequency copyback DI-junctions that were identified matched those isolated by RT/PCR. Our results revealed that defective genomes are generated early during rCDV^RI^ rescue and passage. Independently rescued rCDV^RI^ stocks contained unique defective genomes, out of which one predominant copyback genome persisted alongside the full-length genome during serial passage. We were unable to detect any deletion genomes by RT/PCR, potentially due to their low abundance, as revealed in the Illumina dataset. However, it should be noted that frequency of a DI-junction does not reflect the true quantity of a genome in the sample. Further, as morbillivirus genomes are multiples of six, we expect only rule-of-six-compliant defective genomes as successfully competing with the full-length genome. All high frequency copyback DI genomes that we identified were indeed multiples of six. Whereas recent data on HRSV suggested that DI-genome generation may be sequence-driven rather than a random process^15^, we did not find this to be the case in our studies on rCDV^RI^. Different viruses potentially have different processes for generating/regulating DI genomes, for instance deleting the C protein from some paramyxoviruses increases DI formation ^35–37^.

Next, we combined the use of an attenuated rCDV^RI^ as producer virus with selective UV-inactivation to address DIP production and purification. We successfully demonstrated that such DIPs can be maintained during the course of rCDV^RI^ infection in ferrets. If DIPs are to be used therapeutically then obtaining an appropriate DIP dosage is essential. However, translating dosage from *in vitro* to *in vivo* can be challenging. Our *in vitro* dose-response assay for copyback DIP nDI^RI^04cb determined a DIP to virus ratio at about 10,000:1 (Figure 6B). However, infecting ferrets with a high dose of 70,000 DI genomes:1 virus genome proved inadequate to interfere with virus replication. From our experience using deletion DIP sDI^RI^LedTomTr, we find that DI genome copies do not necessarily translate to infectious DIP titers (unpublished data). Therefore, our true dose may have been well below that required for an *in vivo* system. Our *in vitro* assays using IFN-deficient Vero-cCD150 cells demonstrate that DIPs can inhibit wild-type virus generation. Nevertheless, we believe that any *in vivo* effect using these DIPs or any DIP-based therapy may ultimately be down to innate immune responses mounted by the host.

DIP production is a work in progress and will be an important hurdle to decipher for DIP-based therapeutics. We chose an attenuated producer virus approach for this study. rCDV^RI^ producer virus was based on our previous work demonstrating that introduction of foreign sequences in the second variable hinge of the L protein sufficiently attenuates morbilliviruses ^32,33^. As expected rCDV^RI^Venus(6)LEGFP was also attenuated in ferrets (unpublished data). This is important as such a producer virus would not contribute to disease, thus making it possible to assess a potential interference effect of the DIP. DIPs generated with such a system will be properly packaged and delivered to the right target cells. Additionally, UV-inactivation of such a producer virus would no longer be a safety concern. However, obtaining the high DIP concentrations required using such a system is still a tedious task and any shift in balance results in reduced DIP titers.

DIPs may well have a future as therapeutic interfering particles, but before we can safely make that transition, we need to consider the long-term effects of DIPs, such as their role in viral persistence^38^. Extensive longitudinal experiments in appropriate animal models would need to be carried out to address any safety issues. Although the DIPs in this study were (at our estimated ratio) unable to interfere with wild-type CDV replication, the study provides formal evidence for sustained DI genome replication *in vivo* and provides a valuable basis for future DIP work with (-)RNA viruses.

## Methods

### Cell lines, plasmids and viruses

Vero cells that stably express the canine receptor CD150 (Vero-cCD150) were grown in advanced Dulbecco’s modified Eagle medium (DMEM; Gibco) supplemented with 10% (V/V) fetal bovine serum (HI FBS heat inactivated, Gibco) and GlutaMAX-I (Gibco)^39^. Canine B-cell lymphoma cell line CLBL-1^40^ was grown in RPMI-1640 supplemented with 10% (V/V) FBS. All cells were grown at 37°C and 5% CO_2_.

CDV was isolated from ferret oral swabs and viral consensus sequences were obtained (Accession number KU666057). To construct a plasmid containing the full length genome we obtained seven synthetic fragments with overlapping sequences and assembled them via Gibson Assembly (NEBuilder® HiFi DNA assembly, NEB). A subclone containing the reminder of the viral sequences and restriction sites required to clone in the fragments was generated in a modified pBluescript plasmid^41^. The Gibson assembled fragments were then cloned into the subclone generating pCDV^RI^.

pCDV^RI^ versions expressing reporter proteins Venus, dTom, TagBFP or Gluc were generated by placing the coding sequences as an additional transcription unit at the sixth position of the genome. Briefly, parental plasmid (pCDV^RI^) was linearized using restriction sites *Mse* I at genome position 8340 (H gene) and *Aat* II at genome position 10,696 (L gene), reporter genes (obtained as gBlock Gene Fragments from IDT) were then ligated into the linearized plasmid by Gibson Assembly (NEBuilder® HiFi DNA assembly, NEB). The phosphoprotein gene start and the hemagglutinin (H) glycoprotein gene end sequences were used as signal sequences for the reporter genes. pCDV^RI^Venus(6)-LEGFP was constructed from pCDV^RI^Venus(6). We used restriction site *Avr* II to linearize pCDV^RI^Venus(6) at genome positions 14,141 and 15,060. We then used a 1690 bp synthetic fragment (gBlock Gene Fragments from IDT) containing the digested L gene sequences and EGFP to be inserted by Gibson Assembly (NEBuilder HiFi DNA assembly, NEB). The gBlock contained mutations to replace nucleotides CT at rCDV^RI^Venus(6) genome positions 14,826-27 and 14,835-36 to GG and AA, respectively. These mutations create restriction sites *Msc* I and *Hpa* I in pCDV^RI^Venus(6)-LEGFP which allows EGFP to be swapped easily with other genes when needed.

The expression plasmids encoding N, P and L protein sequences of CDV^RI^ were generated by PCR amplifying the genes from pCDV^RI^ and then ligating them into the eukaryotic expression vector pCG^42^ using *Asc* I and *Spe* I restriction sites to generate pCG-CDV^RI^N, pCG-CDV^RI^P, and pCG-CDV^RI^L.

Similar to the virus rescue plasmids, all defective genome expressing plasmids contained sequences for a T7 promoter, hammerhead ribozyme, defective genome, hepatitis delta virus ribozyme and a T7 terminator. Defective genomes from the T7 transcripts were designed to be in the negative sense orientation. First, a copyback genome core plasmid containing 22 nucleotides of trailer (AGP) and trailer complement sequences with a naturally formed *Eco* RV restriction site was generated. This allowed cassettes (gBlock Gene Fragments from IDT) containing various copyback genomes to be ligated using Gibson Assembly (NEBuilder HiFi DNA assembly, NEB). The deletion genome core plasmid was designed to contain *Eco* RV and *Stu* I restriction sites in the trailer and leader (GP) sequences, respectively. Similar to the copyback plasmids this allows various deletion genome cassettes to be easily ligated using Gibson Assembly. All plasmids described here were sequence verified using Sanger sequencing (Genewiz, USA).

Viruses rCDV^RI^, rCDV^RI^Venus(6), rCDV^RI^dTom(6), rCDV^RI^TagBFP(6), rCDV^RI^Gluc(6) and rCDV^RI^Venus(6)-LEGFP were rescued, grown and titrated in Vero-cCD150 cells by infecting cells with MVA-T7 for 1 h at 37°C. Inoculum was aspirated, and cells were transfected (Lipofectamine 2000, Life Technologies) with pCG-CDV^RI^N, pCG-CDV^RI^P, pCG-CDV^RI^L and pCDV^RI^, pCDV^RI^Venus(6), pCDV^RI^dTom(6), pCDV^RI^TagBFP(6), pCDV^RI^Gluc(6) or rCDV^RI^Venus(6)-LEGFP.

After 18 h the transfection mix was removed and replaced with complete growth medium. Cells were incubated for up to 5 – 7 days at 37°C with 5% (vol/vol) CO_2_. The presence of virus was confirmed by cytopathic effect observed by phase-contrast microscopy and fluorescent microscopy. Virus stocks were grown Vero-cCD150 cells and subjected to one freeze-thaw cycle and debris was removed by centrifugation at 3000 RPM for 10 minutes at 4°C. The cleared supernatant (virus stock) was aliquoted and titrated in Vero cCD150 cells; calculated quantities, expressed in TCID_50_ units were used to calculate M.O.I.s for infections.

### Virus passage

Vero-cCD150 cell monolayers at 2 × 10^5^ cells/ml in T25 flasks were infected with rCDV^RI^Venus(6), rCDV^RI^TagBFP(6), rCDV^RI^dTom(6) or rCDV^RI^Gluc(6) at an MOI of 0.05. At 72 hours post-infection, cells were scraped into culture medium and placed at - 80°C. After freeze-thawing the cells and medium, cell debris were clarified and 2 ml of this was used to infect fresh Vero-cCD150 cells in a T25 flask and so on. Viral titers were determined as TCID_50_/ml for each passage. For the titrations, Vero-cCD150 cell monolayers in 96-well plates were infected with a 10-fold serial dilution of virus sample in OptiMEM. Cytopathic effect (CPE) was recorded about 5 – 7 days post infection and values were calculated by use of the Karber method.

### Copyback genome identification by Sanger sequencing

Total RNA was extracted using TRIzol LS reagent (ThermoFisher) according to manufacturer’s recommendations and RNA pellet resuspended in 40 µl nuclease-free water (Invitrogen). cDNA was generated with 7 µl of resuspended RNA using SuperScript™ III First-strand synthesis system (Thermo Fisher Scientific), and primers A1 and A2 (Table 1) in a total volume of 20 µl. 2 µl of the resultant cDNA was then amplified with primers for copyback genome (A1 and A2, Table 1) and L gene amplification (A1 and B1, Table 1), using Phusion high-fidelity DNA polymerase (NEB) in a total volume of 50 µl (using a touch-down PCR amplification protocol). PCR products were analyzed on a 1% agarose gel and bands gel purified using QIAquick gel extraction kit (Qiagen). Samples were sequenced via Sanger sequencing (Genewiz, USA).

### Defective genome identification by Illumina sequencing

Total RNA was extracted using TRIzol LS reagent (ThermoFisher) according to manufacturer’s recommendations. The RNA pellet was resuspended in 40 µl nuclease-free water (Invitrogen). Library preparation, rRNA depletion and sequencing was carried out according to manufacturer’s recommendations for the MiniSeq system (Illumina). Illumina reads were trimmed of their adaptor sequences and quality checked using an in-house script, and then mapped to the appropriate reference genome with bwa-mem using default parameters (version 0.7.7). Using a modified version^34^ of script chimeric_reads.py v3.6.2^43,44^ chimeric reads were identified amongst each dataset.

### DIP rescue, production and quantification

Vero-cCD150 cell monolayers in 6-well trays were infected with Fowlpox-T7 in Opti-MEM (Gibco) for 30 mins at 37°C and then spinoculated at room temperature for another 30 mins. Medium was then removed and cells transfected with 2 µg of defective genome-expressing plasmid, 0.6 µg of pCG-CDV^RI^-N and 0.4 µg of pCG-CDV^RI^-L for 48 hours. Transfected cells were then superinfected with rCDV^RI^Venus(6)-LEGFP at MOI 0.001. Two days post infection cells were trypsinized and passaged into a T75 flask. Cells were harvested by free-thaw when CPE was visible. rCDV^RI^Venus(6)-LEGFP producer virus was inactivated from all DIP stocks at 60 mJ/cm^2^ in 2 ml culture volume in a 6-well tray using a CX-2000 Crosslinker (UVP).

For quantification, total RNA was extracted using TRIzol LS reagent (ThermoFisher) according to manufacturer’s recommendations and RNA pellet resuspended in 40 µl nuclease-free water (Invitrogen). 3 µl of extracted RNA was used in a TaqMan one-step qRT/PCR assay (Luna^®^ Universal One-Step RT-qPCR Kit, NEB). TaqMan analysis was carried out with primer/probe combination described in Table 1 and analysis performed on the QuantStudio^™^ 6 Flex System (ThermoFisher Scientific).

### DI interference assay

DIP stocks were split in half and UV irradiated to either inactivate the producer virus (60 mJ/cm^2^, active DIPs) or the DIPs (120 mJ/cm^2^ (X8), inactive DIPs). UV irradiation was carried out in 2 ml in a 6-well tray using a CX-2000 Crosslinker (UVP). Vero-cCD150 cell monolayers in 24- or 96-well trays were infected with either active or inactive DIPs for 1 hour at 37°C and then superinfected with rCDV^RI^ for 1 hour at 37°C. Supernatant was then removed and cells washed with phosphate buffered saline (PBS) 3X times. Fresh medium was added onto the cells and incubated for 48 hours. For crystal violet staining, cells were fixed in formalin for 15 mins and then stained with crystal violet for another 15 mins. Cells were washed with water and visualized. For virus titers, cells and medium were freeze-thawed at -80°C and TCID_50_ carried out on Vero-cCD150 cells.

### Animal study design

The animal experiment described here was conducted in compliance with all applicable U.S. Federal policies and regulations and AAALAC International standards for the humane care and use of animals. All protocols were approved by the Boston University institutional animal care and use committee. Twelve 16-week old CDV-seronegative male ferrets (*Mustela putorius furo*) were housed in groups of three. The cages contained toys as a source of environmental enrichment.

Animals were intra-tracheally (IT) inoculated with 10^4^ TCID_50_ rCDV^RI^TagBFP(6). Nine ferrets, divided into groups of three were also inoculated with DIPs. One group was inoculated with DIPs 6 hours prior to rCDV^RI^TagBFP(6). Inoculation was carried out with 2 ml of DIPs or medium and 40 µl rCDV^RI^TagBFP(6) suspension. Animals were monitored several times per day and blood samples were collected every two days post inoculation in Vacuette tubes containing EDTA as an anticoagulant. Procedures were performed under light anesthesia (initial sedation with ketamine, medetomidine and butorphanol followed by maintenance with 1-5% isoflurane in oxygen and atipamazole reversal after handling). All animals were euthanized 14 days post inoculation and necropsies were performed.

### Sample processing and analysis

50 µl of blood sample was analyzed on a VetScan HM5 (Abaxis) to determine total white blood cell (WBC) and lymphocyte counts. Red blood cells (RBC) were then lysed using 1X multi-species RBC lysis buffer (eBioscience). WBC was washed in D-PBS and collected by centrifugation (350g, 10 minutes) and resuspended in 300 – 500 µl D-PBS. The resuspended WBC was used to determine total lymphocyte counts and infected WBC percentages by flow cytometry using a LSRII flow cytometer (BD LSR II, Biosciences). Virus isolation from WBC were performed on Vero-cCD150 cells by titrating the WBC in Opti-MEM.

Ferrets were euthanized by administration of an overdose of barbiturate anesthetic under deep sedation. Lymphoid tissues were collected during necropsy and placed in PBS for single-cell suspensions. Fatty tissue was removed from the lymphoid tissues before dissecting the tissues into smaller pieces to allow dissociation using gentleMACS™ dissociation (C) tubes (Miltenyi Biotec) in Advanced RPMI medium supplemented with 10% fetal calf serum, 1% Glutamax and 1X Antibiotic-Antimycotic (ThermoFisher Scientific). Samples were dissociated on a gentleMACS Dissociator (Miltenyi Biotec) using the m_spleen_C preset parameter. Samples were then passed through cell strainers with 100 µm pore size (Falcon™ Cell Strainers), washed in D-PBS and pelleted by centrifugation (350g, 10 minutes). Pellets were resuspended in an appropriate volume of D-PBS and analyzed by flow cytometry as described above.

## Acknowledgements

We thank Shannon L.M. Whitmer and Jessica R. Harmon at the CDC for their help with NGS and bioinformatic analysis, and Claire Sharp (Tufts University) for providing clinical samples. This work was funded by the Defense Advanced Research Projects Agency (DARPA) INTERfering and Co-Evolving Prevention and Therapy (INTERCEPT) program (HR0011940493) and Boston University.

